# Evolution of VIM-1 producing *Klebsiella pneumoniae* isolates from a hospital outbreak reveals the genetic bases of the loss of the urease-positive identification character

**DOI:** 10.1101/2021.03.02.433680

**Authors:** Nicolas Cabanel, Isabelle Rosinski-Chupin, Adriana Chiarelli, Tatana Botin, Marta Tato, Rafael Canton, Philippe Glaser

## Abstract

Outbreaks of carbapenemase producing *Klebsiella pneumoniae* (CP*Kp*) represent a major threat for hospitals. We molecularly characterized the first outbreak of VIM-1 producing *K. pneumoniae* in Spain, that raised fears about the spread of this strain or of the plasmid carrying *bla*_VIM-1_. Through in-depth genomic analysis of 18 isolates recovered between October 2005 and September 2007, we show that 17 ST39 isolates were clonal, whereas the last isolate had acquired the VIM-1 plasmid from the epidemic clone. The index isolate carried 31 antibiotic resistance genes (ARGs) and was resistant to almost all antibiotics tested. Later isolates further gained mutations in efflux pumps regulators *ramR* and *opxR*, deletion of *mgrB* (colistin resistance) and frameshift mutations in *ompK36* (ß-lactam resistance) likely selected by antibiotic usage. Comparison with publicly available genome sequences and literature review revealed no sign of dissemination of this CP*Kp* strain. However, the VIM-1 plasmid was found in diverse *Enterobacterales* species, although restricted to Spain. One isolate became urease negative following IS*5075* transposition into *ureC*. Analysis of 9755 *K. pneumoniae* genomes showed the same *ureC*::IS*5075* insertion in 14.1% of the isolates and explained why urease activity is a variable identification trait for *K pneumoniae*. Transposition into *ureC* results from the similarity of its 3’-end and the terminal inverted repeats of Tn*21* like transposons, the targets of IS*5075* and related ISs. As these transposons frequently carry ARGs, this might explain the frequent chromosomal invasion by these ISs and *ureC* inactivation in multidrug resistant isolates.

**IMPORTANCE:** Evolution of multidrug resistant bacterial pathogens occurs at multiple scales, in the patient, locally in the hospital or more globally. Some mutations or gene acquisitions, for instance in response to antibiotic treatment, may be restricted to a single patient due to their high fitness cost. However, some events are more general. By analyzing the evolution of a hospital acquired multidrug resistant *K. pneumoniae* strain producing the carbapenemase VIM-1, we showed a likely environmental source in the hospital and identified mutations contributing to a further decrease in antibiotic susceptibility. By combining the genomic analysis of this outbreak with literature data and genome sequences available in databases, we showed that the VIM-1 plasmid has been acquired by different *Enterobacterales* but is only endemic in Spain. We also discovered that urease loss in *K. pneumoniae* results from the specific transposition of an IS element into the *ureC* gene and was more frequent in fluoroquinolone resistant isolates and carrying a carbapenemase gene.

## INTRODUCTION

*Klebsiella pneumoniae* is responsible for a broad range of diseases including pneumonia, blood stream and urinary tract infections, mostly in health-care facilities. *K. pneumoniae* isolates are frequently resistant to multiple antibiotics and contribute to the dissemination of antibiotic resistance genes (ARGs) (1, 2). Carbapenems are among the last resort drugs to treat infections due to multidrug resistant (MDR) *K.pneumoniae* isolates expressing extended spectrum ß-lactamases (ESBL). From the end of the 20^th^ century onwards, the emergence and dissemination of carbapenemase producing *K. pneumoniae* (CP*Kp*) resulting in high mortality rates is becoming a major public health threat. CP*Kp* hospital outbreaks are particularly feared with patient-to-patient transmission or transmission from the hospital environments to patient. Recently, a broad genomic study on CP*Kp* from 244 hospitals in 32 countries across Europe confirmed the existence of dominant lineages responsible for hospital outbreaks (3). In this study, the most prevalent multi locus sequence typing (MLST) types (STs) were from the Clonal Group (CG) 258, including ST258, 512, 340, 437 and 11, expressing the carbapenemase KPC (1, 3). Other prominent CP*Kp* STs are ST307 (4) and ST101 (5). However, the molecular epidemiology of CP*Kp* is different between countries (6) and a large proportion of CP*Kp* isolates belongs to diverse and rare STs denoting relevance of local epidemiology.

In 2007, we reported the first case of a hospital outbreak involving CP*Kp* isolates producing the VIM-1 carbapenemase in a hospital in Madrid, Spain (7, 8). During the same period, *Escherichia coli, Klebsiella oxytoca* and *Enterobacter cloacae* isolates also producing VIM-1 were identified in the same hospital (7). Pulsed field gel electrophoresis (PFGE) of *K. pneumoniae* isolates showed that they were likely clonal (8). This observation raised questions about the risk of endemicity of this clone and of the plasmid carrying *bla*_VIM-1_ (7).

Whole genome sequencing (WGS) is becoming instrumental to decipher hospital outbreaks and to characterize transmission (9). Point mutations and small indels, particularly those leading to gene inactivation or contributing to antibiotic resistance are the main focus of genomic epidemiology studies. Other events, and in particular the mobility of insertion sequences (IS), more difficult to identify by short read sequencing, are frequently set asides. In this work, we have analyzed the evolution of the VIM-1 producing *K. pneumoniae* isolates from the outbreak (7, 8). In addition to mutations selected by antibiotics used in the hospital, we observed a diversity in ARGs and plasmid contents and mobility of transposable elements: a Group 2 intron and three ISs, IS*26*, IS*5075* and IS*421*. In one isolate, IS*5075* transposed into the *ure* operon encoding the urease subunits and led to a urease defective phenotype. By analyzing 9755 publicly available *K. pneumoniae* genome sequences we show that this insertion is frequent, explaining why some *K. pneumoniae* isolates display a urease negative phenotype. Furthermore, through a literature survey and the analysis of publicly available genome sequences, we did not find any evidence of further dissemination of this VIM-1 producing strain. On the other hand, the *bla*_VIM-1_ plasmid has broadly disseminated across *Enterobacterales* species but so far has only been isolated in Spain.

## RESULTS

### Genomic characterization of the outbreak isolates

Illumina WGS of the 18 isolates and *in silico* MLST showed that the 17 first isolates (KP_VIM_1-17) sharing the same PFGE profile belong to ST39 and the last isolate (KP_VIM_18) to ST45 (Table S1). ST45 represents 1.5% (n=161) of the 10,515 genomes retrieved from NCBI (July. 2020). ST39 is less frequent, with only 38 other genome sequences, including seven isolates carrying carbapenemase genes (*bla*_KPC-3_, n=3; *bla*_KPC-2_, n=2; *bla*_NDM-1_, n=2) but none *bla*_VIM-1_. In order to characterize the strain responsible for the outbreak and to identify genetic events occurring during its evolution, we determined the complete genome sequence of the first isolate, KP_VIM_1. KP_VIM_1 chromosome is 5,351,626 base pairs (bp) long. It hosts four plasmids of 227,556 bp (pKP1-1), 110,924 bp (pKP1-2), 76,065 bp (pKP1-3) and 80,027 bp (pKP1-4) (Table S2). The chromosome and plasmids pKP1-1, 2 and 3 carry 31 ARGs (Table S2). Those ARGs target all major classes of antibiotics used against Gram negative bacteria. The porin gene *ompK*35 is interrupted by a non-sense mutation at codon 230. In agreement with the ARG content, KP_VIM_1 is highly resistant to almost all antibiotics tested, remaining susceptible to only fluoroquinolones, tigecycline, and colistin and exhibiting an intermediate phenotype to amikacin, imipenem, meropenem and ertapenem (Fig. S1).

The *bla*_VIM-1_ gene is carried by a gene cassette inserted in a type-1 integron expressing six ARGs in addition to *bla*_VIM-1_ (*aacA4, dfrB1, ant1, cat, emrE* and *sul1*) carried by plasmid pKP1-3 (Fig. 1). BLASTN search using the nucleotide sequence of this plasmid against the contigs of KP_VIM_18 showed 100% identity over its entire length, except a 1722 bp region containing a *catA* gene and missing in KP_VIM_18. The VIM-1 plasmid was therefore likely transferred in the hospital from the outbreak strain to the ST45 *K. pneumoniae* isolate. Plasmid pKP1-3 belongs to IncL/M type. Comparison with complete plasmid sequences showed that pKP1-3 is more than 99.9% identical over 89% of its length to pKP1050-3b carrying *bla*_VIM-1_ from a pan-drug resistant *K. pneumoniae* isolated in June 2016 in a hospital in Madrid (Fig. 1) (10). Both plasmids are highly similar to a *bla*_VIM-1_ carrying plasmid from a *Salmonella* Typhimurium isolated in Spain in 2014 (11) and from *Klebsiella oxytoca* strains isolated in Madrid in 2016 (12). Recently, a closely related plasmid was identified in 28 *Serratia marcescens* VIM-1 producing isolates recovered in our hospital as KP_VIM_1 between September 2016 and December 2018 (13). We identified by BLASTN search ten additional *K. pneumoniae* isolates carrying a plasmid closely related to pKP1-3, among the 85 *K. pneumoniae* genome sequences containing *bla*_VIM-1_ of the 10,515 *K. pneumoniae* genome sequences from the NCBI (Table S3). Strikingly these isolates from four different STs were also all isolated in Spain between 2010 and 2016. Therefore, IncL/M plasmids carrying *bla*_VIM-1_ likely arose in Spain following the insertion of a type 2 integron and disseminated locally only but were recurrently isolated in Spain between 2005 and 2018.

**Fig. 1.**
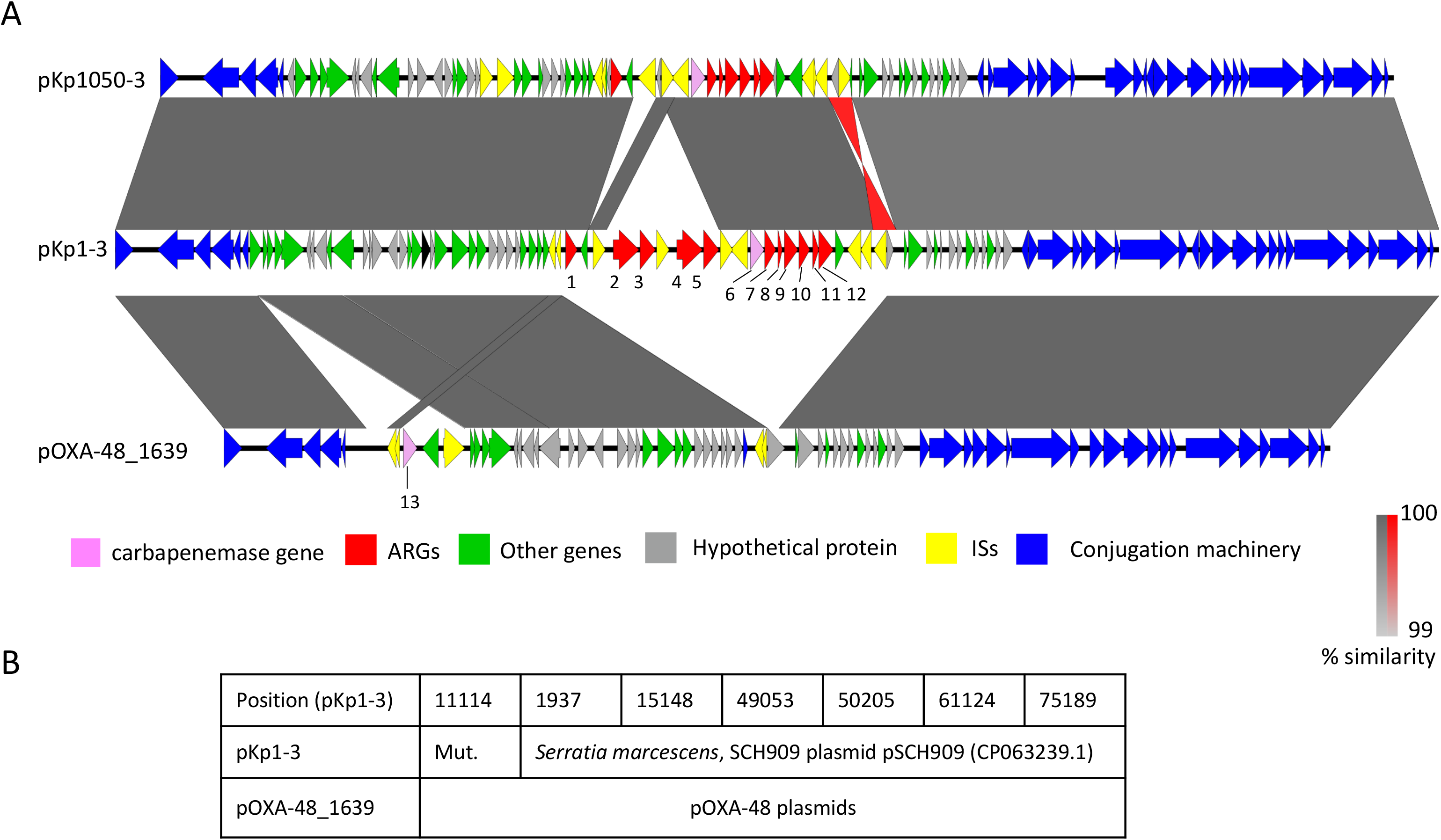
Comparison of pKP1-3, pKp1050-3 and pOXA-48. 1: A. Comparison of plasmids pKp1-3 and pKp1050-3 (Accession: CP023419.1) carrying *bla*_VIM-1_ and of pOXA-48_1639 carrying *bla*_OXA-48_ (Accession: LR025105.1). pOXA48_1639 was chosen as it was the closest relative to pKp1-3. Grey areas between ORFs denote nucleotide identities with a gradient representing 99% (light grey) to 100% (dark grey) identity. In red are represented identities of an inverted region. Genes are indicated by arrows with a color code as in the figure key. Antibiotic resistance genes are numbered as follows, 1: *catA1*; 2 and 4: *msrE_1*; 3 and 5: *mph*E; 6: *bla*_VIM-1_; 7: *aac*A4_2; 8: *dfr*B1 9: *ant* 1_2; 10: *cat* _2; 11: *emr*E; 12: *fol*P_4; 13: *bla*_OXA-48_. B. Analysis of the SNPs detected between pKP1-3 and pOXA-48_1639. Occurrence of SNP among publicly available IncL/M plasmids ware identified by BLASTN. SNPs position in pKp1-3 are indicated in the first line. Mut. indicates that the mutation is specific to IncL/M VIM-1 plasmids. For other positions, plasmids with the pKp1-3 allele or the pOXA-48_1639 allele are indicated in the second and third line respectively. pSCH909 carries *bla*_OXA-10_ and *bla*_TEM-1_, but no carbapenemase gene.

These plasmids are closely related to the broadly distributed IncL/M plasmid pOXA48 carrying the *bla*_OXA-48_ carbapenemase gene (10) (Fig. 1). pKP1-3 shows only seven SNPs over 57,386 conserved bp with pOXA-48_1639, the closest relative identified at the NCBI (accession number LR025105.1). BLASTN search against the NCBI database showed that one SNP was specific to all characterized IncL/M VIM-1 plasmids, whereas for the six other positions, two different allelic forms could be identified: one shared by pOXA-48_1639 and other pOXA-48 plasmids, the other by pKP1-3 and IncL/M plasmids carrying other resistance genes. Therefore, these two plasmids share a very recent common ancestor which acquired either Tn*1999* (14) carrying *bla*_OXA-48_ or an integron carrying *bla*_VIM-1_.

### Intra-hospital evolution of the ST39 lineage follows different paths associated with modifications of antibiotic susceptibility

On the basis of the variants identified, we reconstructed the evolutionary path of the 17 ST39 isolates (Fig. 2A). In total, we identified 64 SNPs (59 in the chromosome and five in the plasmids), and seven short indels, five of which leading to a frameshift in coding frames (Table S4). Ancestral genotype for each polymorphism was predicted by parsimony based on BLASTN comparisons with complete *K. pneumoniae* genomes sequences at the NCBI. The first isolate, KP_VIM_1, shows six SNPs compared to the reconstructed sequence of the last common ancestor (LCA) of the 17 isolates. We next analyzed the root to tip number of chromosomal SNPs according to the time of isolation. Despite the duration of the outbreak over 24 months, we did not observe a strong temporal correlation (Fig. 2B).

**Fig. 2.**
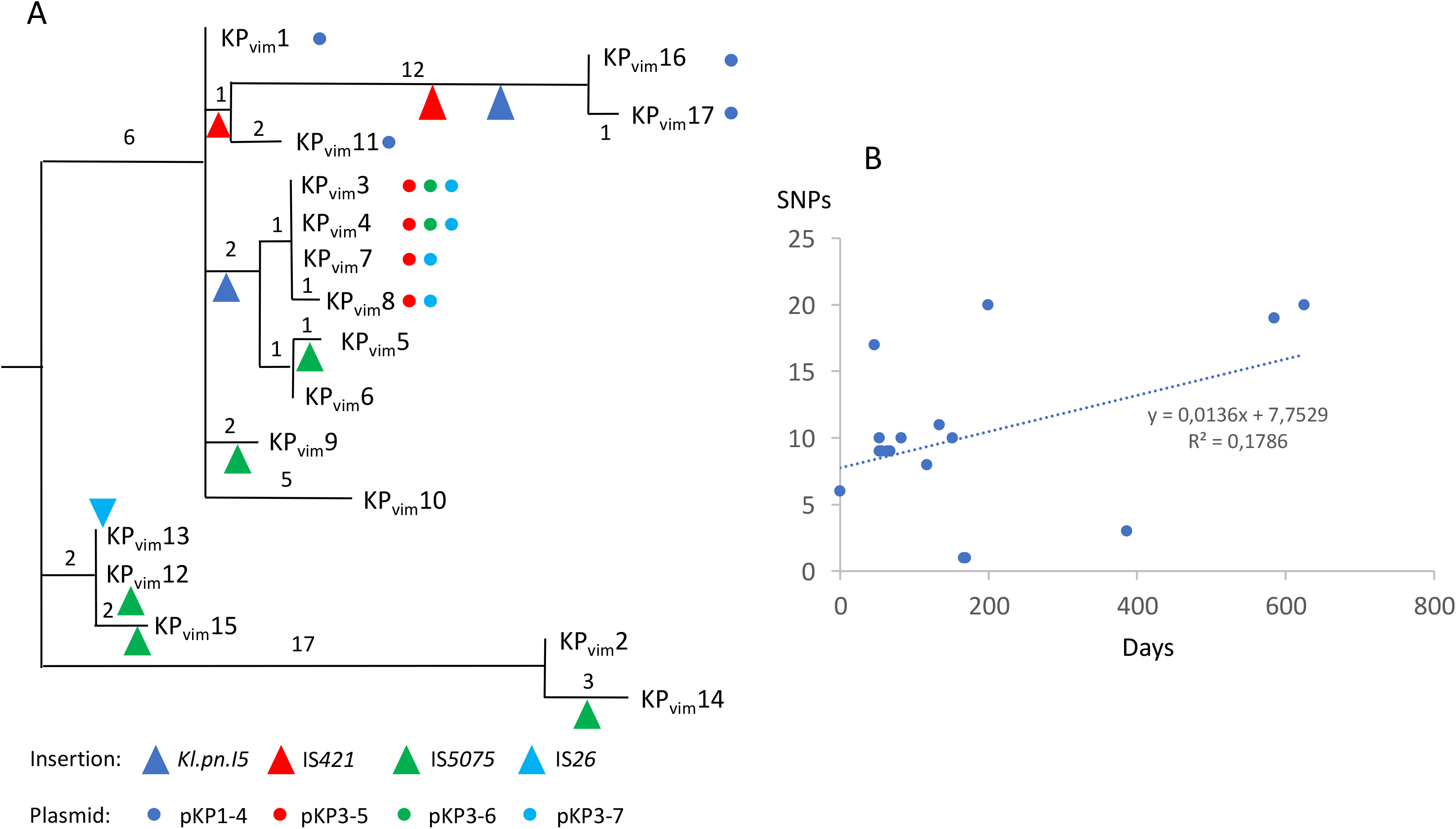
*Hospital evolution of the* K. pneumoniae *ST39 VIM-1 producing strain*. **A.** Phylogeny of the 17 isolates reconstructed by maximum parsimony. Numbers next to branches indicate the number of chromosomal SNPs in the corresponding branch. Presence of plasmids are indicated by colored points and transposition events by triangles. IS*26* insertion in *oqxR* occurred in the common ancestor of KP_VIM_12 and KP_VIM_13. **B.** Root to tip representation of the number of chromosomal SNPs according to the time (in days) following the isolation of the first isolate KP_VIM_1. The trendline equation and the correlation coefficient are indicated on the graph.

We identified three large chromosomal deletions: a 6.3 kb deletion encompassing *mgrB*, a 600 bp deletion of a type 6 secretion system (T6SS) immunity phospholipase A1-binding lipoprotein and a 55.4 kb deletion corresponding to the excision of an Integrated and Conjugative Element. Five large deletions in pKP1-1 and pKP1-2 led to the loss of clusters of ARGs (Table S2 and S4) in agreement with modifications of the antibiotic susceptibility profiles (Table S3).

Several genetic events were likely selected in response to antibiotic use in the hospital. The deletion of the *mgrB* gene led to colistin resistance in KP_VIM_17 (Fig. S1). The same isolate was highly resistant to all ß-lactams including carbapenems due to the inactivation of the second major porin gene, *ompK*36, by a non-sense mutation leading to a stop codon at position 125. In addition, we identified three mutations disrupting *oqxR* and *ramR* genes encoding repressors of efflux systems. *oqxR* was inactivated by an IS*26* insertion in KP_VIM_12 and KP_VIM_13 whereas *ramR* was inactivated by a non-sense mutation in KP_VIM_14 and by a frameshift mutation in KP_VIM_7 and KP_VIM_8. In agreement with previous comparisons of mutants of *oqxR* and *ramR* (15, 16, 17, 18), we observed a stronger decrease in the susceptibility to fluoroquinolones in the isolates mutated in *oqxR* (KP_VIM_12 and KP_VIM_13) and a stronger decrease in tigecycline susceptibility in the isolates mutated in *ramR* (KP_VIM_7, 8 and 14). In the case of KP_VIM_14, the mutation in *ramR* likely compensates the loss of the *qnrA1* gene for fluoroquinolone susceptibility. The five isolates also showed a decreased susceptibility to cefepime and cefoxitin (Fig S2). To assess if there was any fitness cost associated with the increased resistance observed, we followed bacterial growth of these isolates in LB at 37°C. We observed in all four mutated isolates a decreased growth rate compared to KP_VIM_1. The effect was more pronounced for Kp_VIM_17 defective in both *mgrB* and *ompK36* which showed a 17% increase of generation time (Fig. 3).

**Fig. 3.**
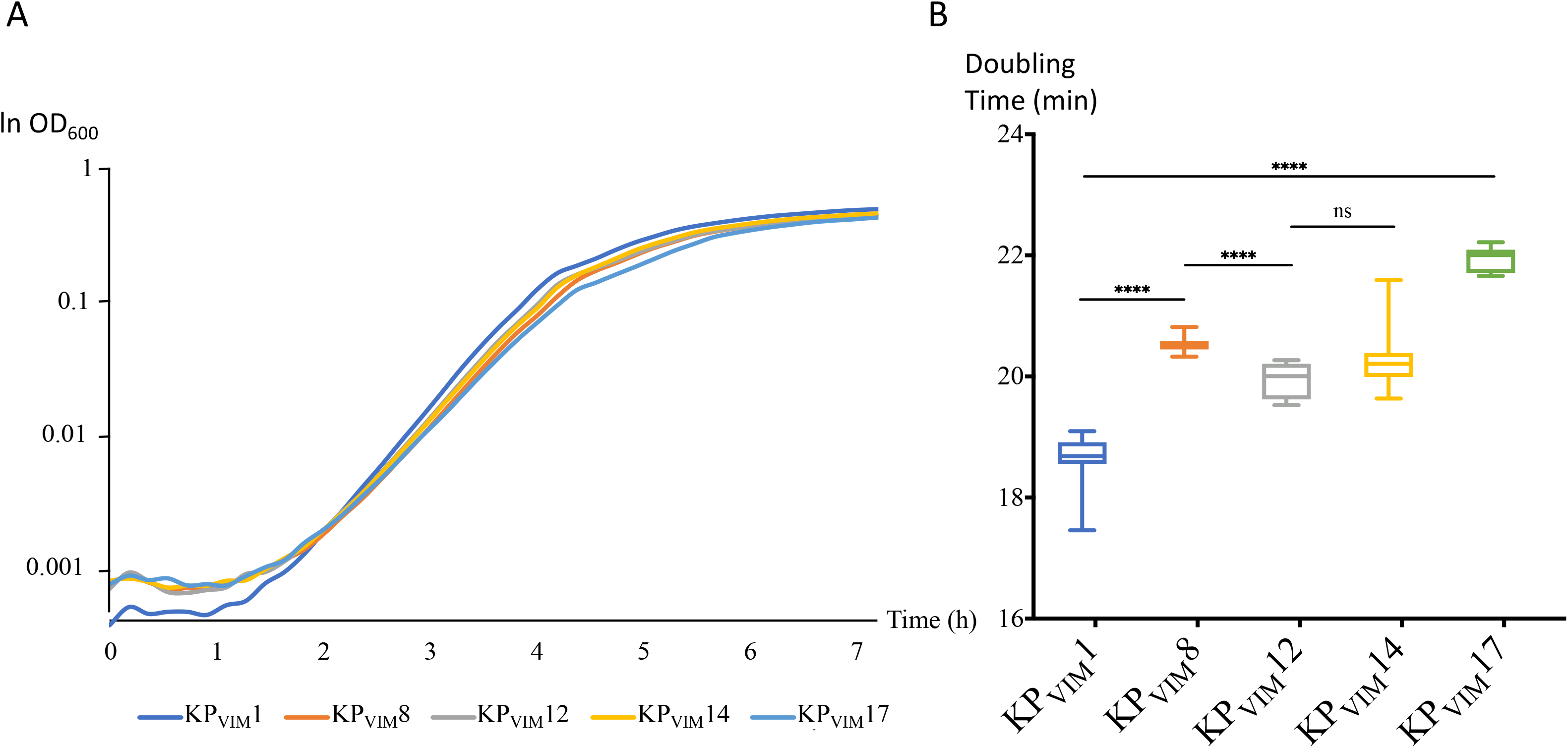
Growth and generation times of isolates with decreased antibiotic susceptibility. **A.** Growth of KP_VIM_1 and of four isolates mutated in a repressor of efflux pumps (KP_VIM_8, KP_VIM_12 and KP_VIM_14) or in *mgrB* and *ompK36* (KP_VIM_17) was followed by using an automatic plate reader. Background was subtracted as described in the Material and methods section. During the first 90 minutes, the OD_600_ was below 0.0015 and its quantification is noisy. B. Box plot representation for 10 replicates of the generation times of the five isolates quantified in early exponential phase 2.5 hours following the start of the culture (OD_600_ between 0.005 and 0.04). Statistical significances were tested with a Student’s t-test. ****, P ≤ 0,0001; n.s. non-significant.

### Diversity of cryptic plasmid content

In the course of the epidemic strain evolution, we also observed changes in plasmid content (Fig. 2). Plasmid pKp1-4 is a IncFII type, which is present in the first isolate KP_VIM_1 and in three of the last isolates of the outbreak (KP_VIM_11, 16, 17), reflecting its stability. This plasmid mainly codes for maintenance functions (toxin antitoxin systems, colicin B production and partition) and conjugative functions. BLASTN search against bacterial genome sequences showed that pKp1-4 is almost identical (99.7% identities over its entire length) to plasmid pEC14III (accession number KU932028.1) from an *E. coli* strain isolated in Finland. We also identified three plasmids specific to the lineage KP_VIM_3 to KP_VIM_8 (Fig. 2). These six isolates share a 34,017 bp-long, linear plasmid (pKP3-5) flanked by two 695 bp-long terminal inverted repeats (TIR). Unlike most linear plasmids described in *K. pneumoniae*, pKP3-5 is unrelated to phages. No adaptive functions were recognized, unlike in a similar linear plasmid pBSSB1 from *Salmonella* Typhi that encodes a flagellin structural gene (19). Search among *K. pneumoniae* genomes revealed 19 isolates carrying putative linear plasmids closely similar to pKP3-5 (>90% identities over 90% of the length). The two other plasmids are small high copy number plasmids: pKP3-6 (2811 bp) and pKP3-7 (3861 bp) that are present in strains KP_VIM_3 to 6 and KP_VIM_3 to 8 respectively (table S2 and Fig. 2). No adaptive functions were predicted in these two plasmids. For these three plasmids, we could not determine whether they were gained in the common ancestor of the KP_VIM_3 to KP_VIM_8 clade or lost by other isolates.

### Insertion of IS*5075* into *ureC* is responsible for a urease negative phenotype in one isolate of the outbreak

In addition to IS*26* insertion in *oqxR*, we identified nine transpositions of mobile genetic elements: two insertions of a class 2 intron named *Kl.pn.I5* (20), and two and five transpositions of IS*421* and IS*5075* respectively (Fig. 2). Compared to the other isolates, KP_VIM_14 was characterized by an IS*5075* inserted three codons upstream of the stop codon of the *ureC* gene encoding the urease catalytic subunit (Fig. 4A). This insertion led to a *ureC* - IS*5075* transposase gene fusion. It might also have a polar effect on the expression of the downstream genes of the operon: *ureE*, *ureF* and *ureG*. Accordingly, the KP_VIM_14 isolate was urease negative, whereas all other isolates from the outbreak were urease positive (Fig. 4B). IS*5075*, like its close relative IS*4321*, is known to transpose into the TIR of Tn*21* and of related transposons of the Tn*3* family (21). Tn*3* family transposons are abundant and diverse (22). They are vectors of heavy metal resistance and ARGs (21). The 17 ST39 isolates harbor three copies of IS*5075* inserted in a pKP1-2 Tn*3* family transposon, just after the initiation codon of a pKP1-1 gene coding for a EAL motif protein and upstream a chromosomal permease gene (Fig 4A). Four independent and identical transposition events of IS*5075* also occurred in the TIR of a Tn*3* family transposon carried by pKP1-3, in KP_VIM_5, KP_VIM_9, KP_VIM_12 and KP_VIM_15 (Fig. 2A). Based on the conservation of the insertion sites of IS*5075* we proposed a 13 bp consensus sequence for the IS*5075* transposition site (Fig. 4A).

**Fig. 4.**
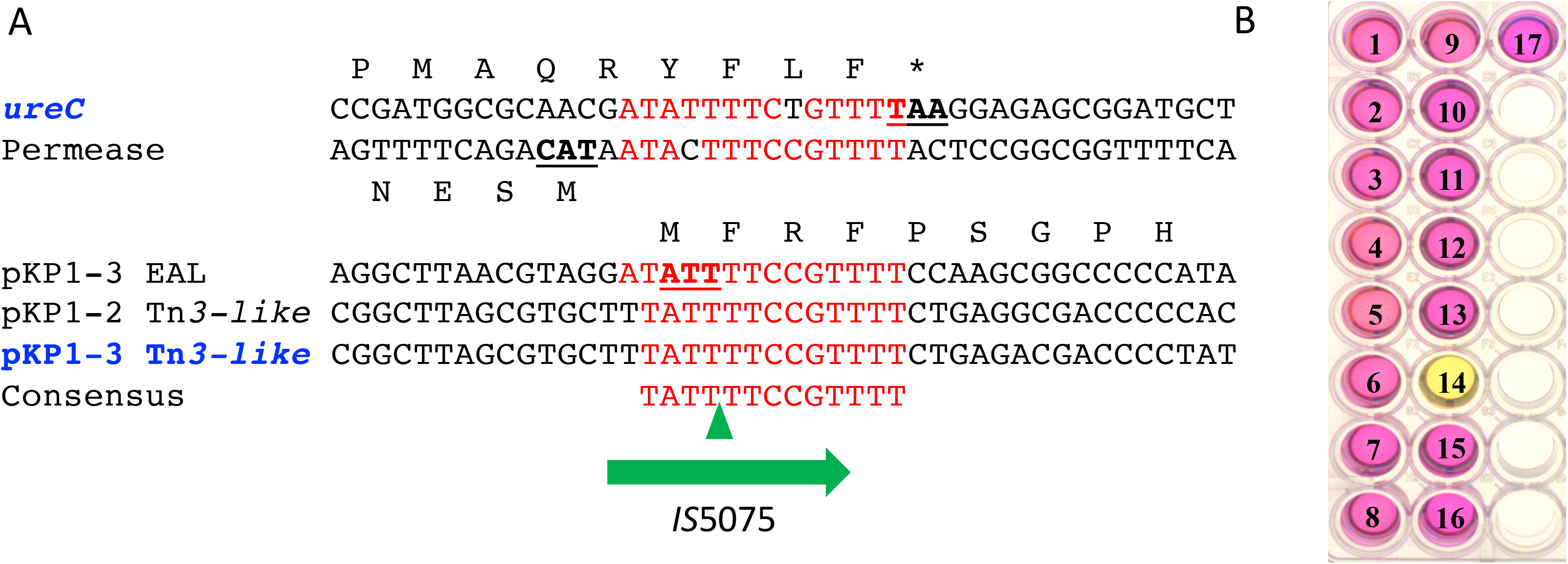
Urease inactivation following IS5075 transposition. **A.** Sequence alignment of the sites targeted by IS*5075* among KP_VIM_ isolates. In blue, targets of transposition events occurring during the outbreak: *ureC* in KP_VIM_14 and pKP1-3 Tn*21* in KP_VIM_5, KP_VIM_9, KP_VIM_12 and KP_VIM_15. The green triangles correspond to IS*5075* insertion sites. In red are indicated conserved bases. Stop and start codons are underlined. B. Urease activity test of the 17 ST39 isolates. The number of each KP_VIM_ isolate is indicated on the well. A pink color of the indole reaction reveals a urease positive phenotype.

### Urease negative phenotypes are prevailing in several *K. pneumoniae* MDR lineages

Urea hydrolysis is an identification trait of *K. pneumoniae* in clinical microbiology laboratories. However, earlier reports have shown that 5% of *K. pneumoniae* isolates are urease negative (23). In order to determine whether this phenotype was due to similar IS*5075* transposition, we analyzed the *ureC* gene in 9755 *K. pneumoniae* genomes quality filtered from the 10,515 genome sequences retrieved from the NCBI (Table S5). BLASTN search showed that an IS*5075* or a similar IS was inserted at the same position in 1380 isolates (14.1%) (Table 1). Search for other insertions or frameshifts in *ureC* did not reveal other frequent mutations putatively responsible for a urease deficiency.

**Table 1:** Comparison of *ureC*::IS*5075* and *ureC* WT *K. pneumoniae* isolates for antibiotic resistance features and ARG and IS copy numbers.

|  |  | All | *ureC* WT | *ureC*::IS5075 | <sup>§</sup>P values |
| --- | --- | --- | --- | --- | --- |
| #Number of isolates | All isolates | #9755 | 8375 | 1380 <sup>@</sup>(14.1%) |  |
|  | Minus ST11 ST14 | 7978 | 7540 | 438 <sup>@</sup>(5.4%) |  |
| *gyrA* or *parC* QRDR mutated | All isolates | 6062 | 4763 <sup>§</sup>(55.9%) | 1299 <sup>&</sup>(94%) | 1e-153 |
|  | Minus ST11 ST14 | 4367 | 4009 <sup>§</sup>(53.2%) | 358 <sup>&</sup>(81.7%) | 3e-31 |
| Carbapenemase gene | All isolates | 5146 | 3953 <sup>§</sup>(47.2%) | 1193 <sup>&</sup>(86.4%) | 6.3e-161 |
|  | Minus ST11 ST14 | 3677 | 3393 <sup>§</sup>(45%) | 284 <sup>&</sup>(64.8%) | 8.4e-16 |
| Carbapenemase gene and *gyrA* or *parC* mutated | All isolates | 4549 | 3376 <sup>§</sup>(40.3%) | 1173 <sup>&</sup>(85%) | 2.2e-208 |
|  | Minus ST11 ST14 | 3093 | 2829 <sup>§</sup>(36.5%) | 264 <sup>&</sup>(60.2%) | 3.3e-21 |
| Average number of IS5075 and related IS | All isolates | 1.82 | 1.31 | 5 | 0 |
|  | Minus ST11 ST14 | 1.45 | 1.27 | 5,11 | 4.2e-207 |
| Average ARGs number | All isolates | 9.33 | 8.81 | 12.5 | 4.3e-101 |
|  | Minus ST11 ST14 | 8.7 | 8.51 | 11.74 | 1.6e-23 |

<sup>#</sup>After filtering out 760 genome sequences out of the 10,515 sequences retrieved from the NCBI. <sup>@</sup>% of isolates with an IS insertion in *ureC*. <sup>§</sup>% of *ureC* WT isolates mutated in QRDR and/or carrier of carbapenemase genes. <sup>&</sup>% of *ureC*::IS5075 isolates mutated in QRDR and/or carrier of carbapenemase genes. <sup>§</sup>Significance of the difference between *ureC*::IS5075 and *ureC* as determined by the Chi-square or the Wilcoxon Rank sum statistical tests.

To determine whether the insertion of IS*5075* into *ureC* preferentially occurred under specific genetic backgrounds, we analyzed the 45 *K. pneumoniae* STs with at least 20 isolates (Fig. 5). We observed that IS*5075* urease inactivation occurred throughout the species with variable frequencies. In seven STs, all with less than 100 isolates, no insertion was observed. On the other hand, we observed a high proportion of *ureC*::IS*5075* isolates in some STs like ST11 (884 out of 1603) and ST340 (18 out of 77 isolates) from the clonal group (CG) 258 and ST14 (58 out of 174). On the other hand, the two-other dominant CG258 STs, ST258 and ST512, showed lower insertion frequencies of 6.9% and 4.1% respectively.

**Fig. 5.**
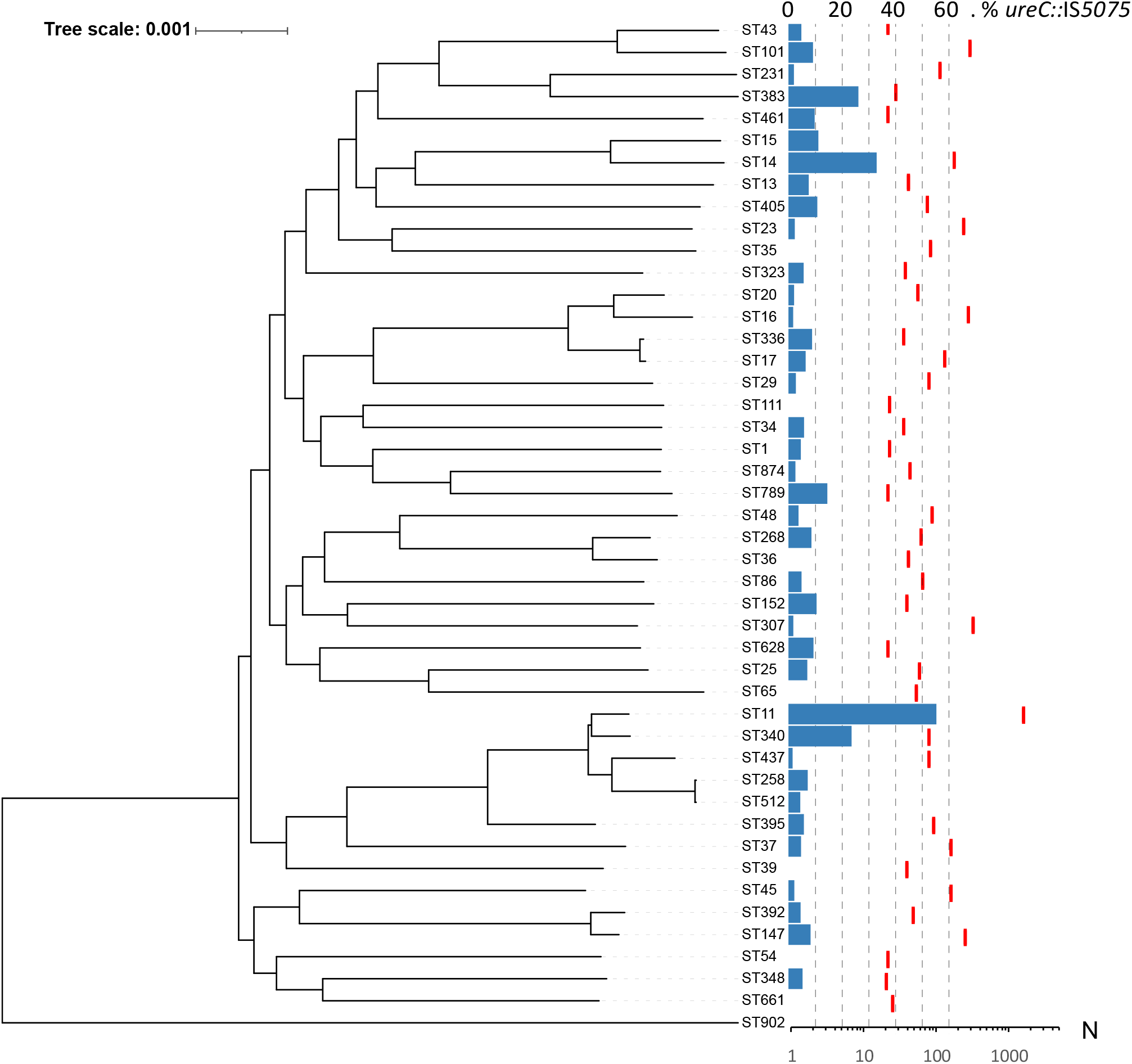
*Distribution of IS5075 insertions in ureC among* K. pneumoniae *isolates*. Occurrence of IS*5075* insertion among the 45 STs with at least 20 isolates among 9755 *K. pneumoniae* genome sequences retrieved from the NCBI. Phylogeny was reconstructed using Parsnp (50) and by using a representative isolate from each ST. The tree was rooted according to David et al. (3). Blue bars indicate the % of isolates with an insertion in *ureC* (upper scale) and red dashes the number of isolates in the corresponding ST (lower scale)

As several of the STs associated with a higher frequency of *ureC*::IS*5075* include major MDR lineages, we next analyzed the distribution of IS insertions in *ureC* in relation with antibiotic resistance. As markers of antibiotic resistance, we considered mutations in fluoroquinolone resistance (FQR) determinants, presence of carbapenemase genes and the number of ARGs among the 9755 *K. pneumoniae* genomes sequences (Table S5). Among these genome sequences, 62% were mutated in *gyrA* and/or *parC* quinolone resistance-determining regions (QRDR), 53% carried carbapenemase genes, and the average number of ARG was 9.33, revealing a strong bias towards MDR isolates (Table 1). Despite this bias, *ureC*::IS*5075* isolates appeared as even more resistant, with an average number of 12.5 ARGs compared to 8.8 in the remaining isolates, 94% of the isolates showing mutations in *gyrA* and/or *parC* and 86.4% carrying a carbapenemase gene (Table 1). To determine whether the insertion in *ureC* was associated with a global expansion of IS*5075* and related ISs, we estimated the copy number of these ISs in the different isolates (Table 1). Isolates with an IS insertion in *ureC* showed in average a 4-fold higher copy number of IS*5075* and related ISs than the remaining isolates (5 *vs*. 1.31). On the other hand, more than half of the isolates with an intact *ureC* genes did not carry a single IS*5075* copy (4334 out of 8375).

In a given ST a high frequency of *ureC*::IS*5075* isolates might result from frequent transposition events or from the expansion of lineages carrying the insertion. To discriminate between these two possibilities, we performed a whole genome phylogeny focusing on ST11, ST14 and ST258. ST11 was the most abundant ST among the genome sequences retrieved from the NCBI (16.4% of all isolates). Except two isolates WT for *gyrA* and *parC*, all ST11 isolates were predicted to be FQR (Fig. 6). The two most populated lineages belong to the K-types KL64 (n=622) and KL47 (n=463). These closely related lineages share the same three mutations in QRDR regions (ParC-80I, GyrA-83I, GyrA-87G) and carry the carbapenemase gene *bla*_KPC-2_. Analysis of IS*5075* insertions in *ureC* showed an uneven distribution, mostly associated with these two lineages. In the KL64 clade, the IS insertion is ancestral, as it was present in all except six isolates (in pink). In the KL47 clade, two different situations were noted: an ancestral transposition event in the LCA of a specific sublineage, with the 138 isolates from this clade showing an IS*5075* in *ureC* (clade colored in red); a relatively high frequency of insertion in the other isolates of the clade (85 out of 324, 26%) likely resulting from multiple sporadic transposition events. Out of the two clades, the frequency of insertion is much lower (8.5%). All over the ST11 phylogeny, insertion in the *ureC* gene was associated with a higher copy number of IS*5075* with on average 5 copies compared to 1.7 in ST11 isolates with a WT *ureC* gene. Altogether these results show that the high proportion of ST11 isolates mutated in *ureC* results in a large part from the dissemination of two clades showing a high number of IS*5075* copies. The situation was similar among ST14 isolates, as all but one isolate (n=58) mutated in *ureC* belonged to a single FQR lineage suggesting that transposition occurred in the LCA of the lineage (in blue, Fig. S2). Isolates of this lineage also showed a high IS*5075* copy-number (n=5.1). In ST258, isolates were characterized by a lower frequency of IS insertion in *ureC* (6.9%). Most of the ST258 isolates cluster in two lineages expressing two different capsule operons of K-type KL106 and KL107 and associated mostly with the carbapenemase genes *bla*_KPC-2_ and *bla*_KPC-3_ respectively (24). In contrast to what was observed in ST11 and ST14, no expansion of a large *ureC*::IS*5075* clade occurred (Fig. S3). All but two isolates with the insertion in *ureC* belonged to the KL107 lineage. Strikingly, this clade was characterized by an higher copy number of IS*5075* of 2.17 (5.24 for *ureC*::IS*5075* isolates) compared to only 0.12 for the KL106 lineage. Therefore, a major driver for insertion into *ureC* is the presence of an IS*5075* or a related IS and its active transposition.

**Fig. 6:**
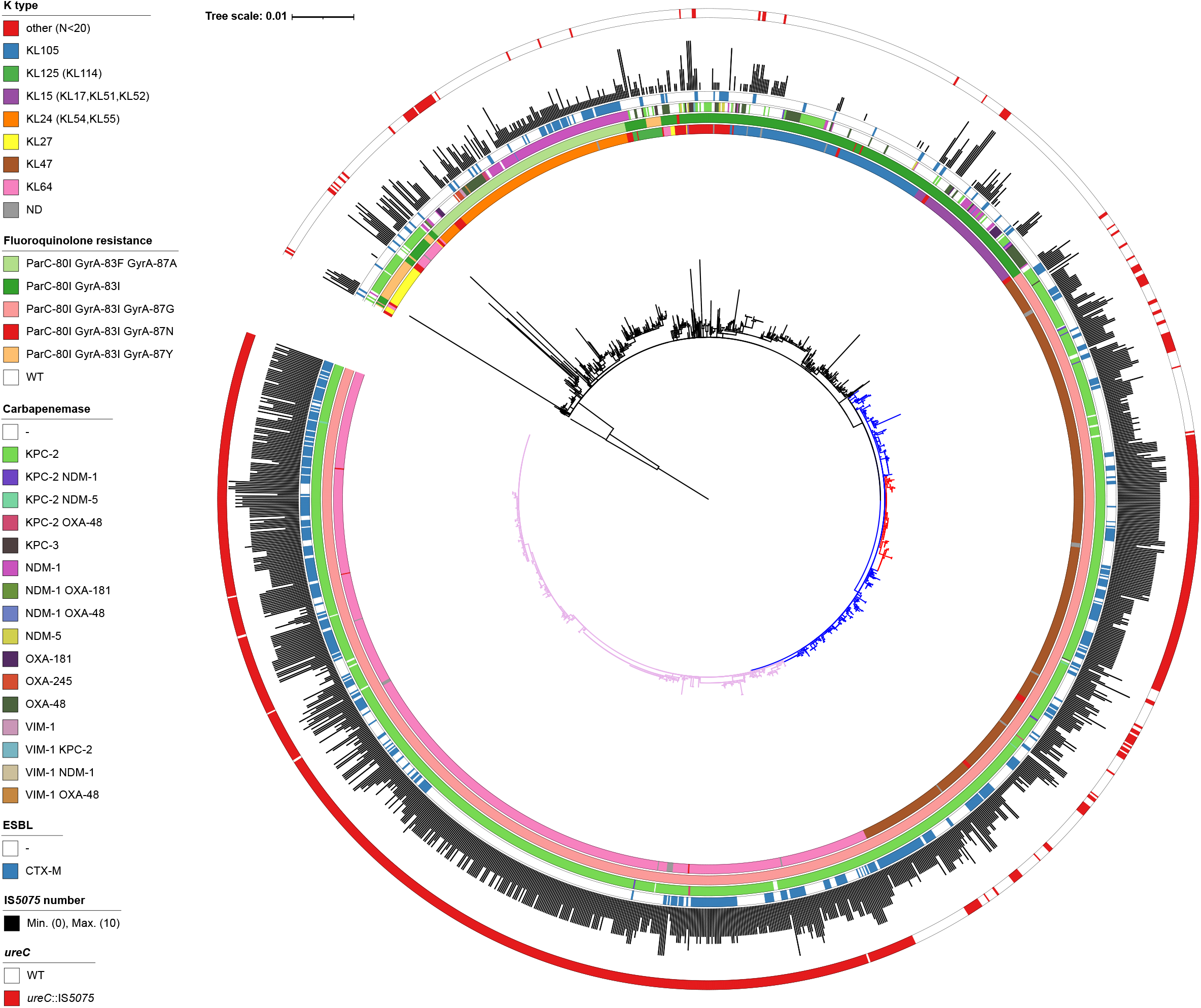
*Core genome phylogeny of* K. pneumoniae *ST11 isolates*. Phylogeny was obtained by using Parsnp (50) considering 1603 genomes passing the quality threshold. K-type, mutations in *gyrA* and *parC* QRDR, carbapenemase genes, *bla*_CTX-M_ genes, copy-number of IS*5075* and related ISs and IS insertion in *ureC* are annotated by circles from inside to outside as indicated in the figure key (left). The *ureC* deficient KL64 lineage is in pink. The KL47 lineage is in blue and the *ureC* deficient sublineage in red. The two *gyrA/parC* WT isolates were used as outgroups to root the tree. The tree was visualized by using iTOL (51).

## DISCUSSION

Whole genome sequencing has revolutionized molecular epidemiology and its use in outbreak analysis has contributed to decipher the path of pathogen transmission (25). Here, we investigated the first outbreak due to a VIM-1 producing *K. pneumoniae* in Spain (7, 8). The strain was extensively drug resistant and belongs to an uncommon ST (ST39). Based on available genomic data, we showed that the strain pre-existed in the hospital prior to the identification of the first isolate in October 2005. Furthermore, the weak temporal signal in the evolution (Fig. 2B) indicated a likely environmental reservoir in the hospital, which agrees with epidemiological data (7). Molecular clock for *K. pneumoniae* evolution have been estimated between 1.4 (26), 1.9 (27) and 3.65 (28) mutations/10^6^bp/year. Here, the rate of SNPs/10^6^bp/year is on the lower range (n=0.87). Growth as a biofilm compared to planktonic growth has been related to a greater diversity due to its structured organization, but a lower mutation rate due to a reduced number of generations (29). The diversity observed, the duration of the outbreak and the small number of SNPs agree with a biofilm source of the isolates. In line with this observation, we observed biofilm production of all the isolates but to variable levels (Fig. S4).

During the 2-year evolution of the strain, we observed variations in the antibiotic resistance profile. This was due on the one hand to the loss of ARGs (Table S4). On the other hand, mutations leading to the increased expression of efflux pumps or to a decreased drug permeation, and subsequently to a decreased susceptibility to some antibiotics, were selected. However, these mutations led to a fitness cost (Fig. 3), which might explain their limited expansion in the hospital.

By combining genomic analysis of the strain responsible for the outbreak with global genomic information retrieved from the NCBI and data from the literature, we were able to draw more general conclusions related to the risk associated with the outbreak strain and the VIM-1 plasmid. Likewise, we were able to identify the main reason for urease deficiency among *K. pneumoniae* isolates. Following the identification of the first VIM-1 isolates in Spain, their dissemination was a matter of concern (7). Although we showed that one single ST39 clone, except for one isolate, was responsible for the outbreak, we did not identify any new occurrence of this strain or of a ST39 isolate carrying *bla*_VIM-1_ based on bibliographical survey and on the analysis of more than 10,000 *K. pneumoniae* genome sequences publicly available. Therefore, this clone seems to be restricted to the hospital where it was isolated. Conversely, we showed that the plasmid carrying *bla*_VIM-1_ has disseminated among various *Enterobacterales* species. Transfers occurred probably in the hospital context, as suggested in the case of a *S. typhimurium* isolate (11). Similarly, we showed the transmission of the VIM-1 plasmid between *K. pneumoniae* isolates in the course of the outbreak. We previously predicted similar transfers between *K. pneumoniae* and *E. coli* based on plasmid typing and size determination (7). This IncL/M plasmid is closely related to the broadly disseminated pOXA48. Our mutation analysis strongly suggests independent gain of a carbapenemase gene by very similar plasmid backbones showing only seven SNPs over 57,386 bp. In agreement with this hypothesis, the first OXA-48 plasmid was detected in Spain in 2009 (30) four years after the first VIM-1 isolate of the hospital outbreak (7).

Strikingly, IncL/M VIM-1 plasmids were until now only reported in Spain. A recent study on plasmids encoding VIM-1 from broad origins showed that among the 28 plasmids analyzed, nine were from IncL/M type (31). These nine plasmids were related to pKP1-3 and were from *K. pneumoniae*, *E. hormaechei* and *E. cloacae* and all from Spain. The limited dissemination of the VIM-1 plasmid might be due to the conjunction of different factors including: a lower conjugation efficiency than pOXA-48 plasmids, a fitness cost restricting its dissemination to environments characterized by strong selective pressures, such as the hospital, or a specificity in antibiotic prescription in Spain. Comparing IncL/M VIM-1 and OXA-48 plasmids provides a model system to study two closely related plasmids with two different spreading destinies.

Urease is considered in many bacterial species as a virulence factor beyond its contribution in harnessing urea as a nitrogen source (32). Urease participates in the adaptation to acidic conditions in a broad range of human pathogens, including *Helicobacter pylori* (33), *Yersinia enterocolytica* (34) and *Proteus mirabilis* (35). Urease is considered as a potential target for the development of new antibacterial drugs against enteric bacteria including *K. pneumoniae* (36). In *K. pneumoniae*, the urease has been shown to contribute to gastrointestinal colonization (37). However, a significant proportion of *K. pneumoniae* isolates are urease negative. Here, we showed that the inactivation of the operon is due to the transposition into the *ureC* gene of IS*5075* or of related ISs, like IS*4321* sharing the same specificity. Urease inactivation can be observed in both carriage isolates and isolates associated with clinical symptoms. For instance, we identified a cluster of eight IS*5075*:: *ureC* ST340 isolates from a single institution (Fig. S5). These isolates were recovered from three patients, from urinary tract infections, blood culture, cerebrospinal fluid and fecal carriage (38).

Among *ureC*::IS*5075* isolates, we observed a higher prevalence of *gyrA* and *parC* mutations and of carbapenemase genes and more generally, a higher number of ARGs compared to *ureC* WT isolates (Table 1). This was partly due to a small number of MDR lineages mutated in *ureC*, such as those of ST11 and ST14, which represent 68% of the *ureC*::IS*5075* isolates (Fig 6 and Fig. S2). Nevertheless, this higher prevalence remained true even after removing ST11 and ST14 isolates (Table 1). IS insertions in *ureC* were also associated with a four-fold increase in IS*5075* copies, resulting from additional transposition events (Table 1). This expansion of IS*5075* in some genetic backgrounds might be a relatively recent event. Indeed, 44% of the isolates did not carry a single IS*5075* copy, despite the high number of ARGs in the genomes we have analyzed. Indeed, IS*5075* most frequent targets are the conserved TIR of transposons related to Tn*21*, which are ARG vectors and frequently carried by conjugative plasmids as in the case of pKP1-2 (21, 22). This insertion specificity represents a safe harbor for these ISs, as it does not incur fitness costs and ensures their dissemination. The insertion into *ureC* results from the high similarity between its last codons and TIRs of Tn*21* and is likely accidental. Therefore, the higher frequency of *ureC* inactivation in some MDR lineages might merely be a consequence of a more frequent acquisition of Tn*3* family transposons carrying IS*5075*. However, we cannot completely dismiss the possibility that the loss of urease activity might provide MDR *K. pneumoniae* isolates with a selective advantage under some circumstances. This seems rather unlikely, as other *ureC* inactivation events, including transpositions of other IS, would have been expected in that case and we did not detect such events. Overall, IS*5075* transposition into *K. pneumoniae ureC* gene represents a perfect example of chromosomal colonization by IS elements carried by plasmids and leading to a homoplasic loss of function.

## Material and methods

### Bacterial strains, growth conditions and antibiotic susceptibility testing

VIM-1-producing *K. pneumoniae* isolates were collected from 2005 through 2008 at Ramon y Cajal University Hospital in Madrid, Spain (8) (tableS1). Colistin Minimum Inhibitory Concentration (MIC) was determined in Mueller Hinton (MH) broth as recommended by the Clinical & Laboratory Standards Institute guidelines (CLSI) (39). Susceptibility against 33 other antibiotics (Fig. S1) was evaluated by disk diffusion on MH agar according to the CLSI guidelines (39). Fitness was determined by growth curve analysis with an automatic spectrophotometer Tecan Infinite M200 during 24 hours in LB. Wells were inoculated with overnight cultures at an OD_600_ of 0.001. OD_600_ was measured every ten minutes. Background was determined as the average value of the OD_600_ of the three first time points. Doubling time was determined between OD_600_ 0.005 and 0.03, where an almost perfect fit with an exponential growth was observed.

### Genome sequencing and sequences analysis

*K,pneumoniae* genomes were sequenced by using the Illumina HiSeq2500 platform, with 100-nucleotides paired-end reads. Libraries were constructed by using the Nextera XT kit (Illumina). Reads were assembled with SPAdes 3.9.0 (40). The complete genome sequence of strain KP_VIM_1 was determined by using the long-read PacBio technology (Macrogen, Seoul, Korea). Reads were assembled with the RS_HGAP_Assembly.3 protocol (41) and with Canu (42). The consensus sequence was polished with Quiver (41) and manually corrected by mapping Illumina reads with Breseq 0.33.2 (43). Variants compared to KP_VIM_1 were identified by using Breseq (43). Genome sequences were annotated with Prokka 1.14.5 (44) and analyzed for MLST and ARG content by using Kleborate (45) and Resfinder 4.0.1 (46). Plasmid incompatibility groups were identified by using PlasmidFinder 2.1 (47). Directionality of mutations was determined as previously described by performing BLASTN comparisons against publicly available *K. pneumoniae* genomes (48).

### Analysis of publicly available genome sequences

*K. pneumoniae* genome assemblies (n=10,515) were downloaded from the NCBI (July 2020) with Batch Entrez (49). Genome sequences with more than 200 contigs of more than 500 nt were filtered out. Sixty genome sequences (Bioproject PRJNA510003) for which the contig ends corresponding to repeated sequences have been trimmed were removed from the analysis. In total, we analyzed IS*5075* insertions in 9755 genome sequences (Table S5). Phylogenetic analysis was performed by using Parsnp 1.1.2 (50). Recombination regions were visually identified as regions with a higher SNP density by using Gingr (50) and removed from the reference genome sequence (ST11: strain FDAARGOS_444, CP023941.1; ST14: strain 11, CP016923.1; ST258: strain BIC-1, NZ_CP022573.1; ST340: strain EuSCAPE_RS081, GCA_902155965.1_18858_1_51). Insertion of IS*5075* and of related ISs in *ureC* was identified by BLASTN search using as query sequence the junction sequence detected in the KP_VIM_14 isolate encompassing 20 nt of the *ureC* gene and 20 nt of IS*5075* (E-value of 1e^−10^ as threshold). The integrity of *ureC* was tested by tBLASTN using the UreC protein sequence from KP_VIM_1 as query. Copy number of IS*5075* and of closely related ISs was estimated by counting BLASTN hits (100% identity over the entire length), using the first 17 nucleotides of IS*5075* sequence as query. Phylogenetic trees were visualized by using iTOL (51).

### Phenotypic analyses

Urease detection test was carried out with urea-indole medium (BIORAD) according to the manufacturer’s instructions. Biofilm formation capacity was measured by the microtiter plate assay as previously described (52). *K. pneumoniae* strain LM21 (53) was used as a positive control.

### Statistical analysis

The significance of the differences in frequencies of IS insertions in *ureC* was determined by using the Chi-square test. The significance of differences in IS*5075* copy numbers and in ARGs numbers was determined by the Wilcoxon Rank sum tests. Both tests were performed by using standard libraries contained within the R statistics package (http://www.R-project.org/). Statistical significances of growth rate differences were tested with a Student’s t-test.

## Availability of data

All sequence data have been deposited at DDBJ/EMBL/GenBank (Bioproject PRJEB41835) with the following accession numbers: LR991401, KP_VIM_1 chromosome and plasmids; LR991487, plasmid pKP1-5, LR991544, plasmid pKP1-6; LR991565, plasmid pKP1-7. Biosamples for the Illumina sequence data are listed in Table S1.

## Acknowledgments

This work was supported by grants from the French National Research Agency (ANR-10-LABX-62-IBEID), and from the European Union’s Horizon 2020 Research and Innovation Program under Grant Agreement No. 773830 (Project MedVetKlebs, One Health EJP). Adriana Chiarelli is part of the Pasteur - Paris University (PPU) International PhD Program, with funding from the Institut Carnot Pasteur Microbes & Santé, and the European Union’s Horizon 2020 research and innovation programme under the Marie Sklodowska-Curie grant agreement No 665807.

The authors thank Rafael Patiño-Navarrete for his help in the bioinformatics analysis and Laurence Ma from the Institut Pasteur Biomics platform for her help in Illumina sequencing.

